# High Peak Density Artifacts in Fourier Transform Mass Spectra and their Effects on Data Analysis

**DOI:** 10.1101/191205

**Authors:** Joshua M. Mitchell, Robert M. Flight, Qing Jun Wang, Woo-Young Kang, Richard M Higashi, Teresa W-M Fan, Andrew N. Lane, Hunter N.B. Moseley

## Abstract

Fourier-transform mass spectrometry (FT-MS) allows for the high-throughput and high-resolution detection of thousands of metabolites. Observed spectral features (peaks) that are not isotopologues do not directly correspond to known compounds and cannot be placed into existing metabolic networks. Spectral artifacts account for many of these unidentified peaks, and misassignments made to these artifact peaks can create large interpretative errors. Without accurate identification of artifactual features and correct assignment of real features, discerning their roles within living systems is effectively impossible.

We have observed three types of artifacts unique to FT-MS that often result in regions of abnormally high peak density (HPD), which we collectively refer to as HPD artifacts: i) fuzzy sites representing small regions of m/z space with a ‘fuzzy’ appearance due to the extremely high number of peaks present; ii) ringing due to a very intense peak producing side bands of decreasing intensity that are symmetrically distributed around the main peak; and iii) partial ringing where only a subset of the side bands are observed for an intense peak. Fuzzy sites and partial ringing appear to be novel artifacts previously unreported in the literature and we hypothesize that all three artifact types derive from Fourier transformation-based issues. In some spectra, these artifacts account for roughly a third of the peaks present in the given spectrum. We have developed a set of tools to detect these artifacts and approaches to mitigate their effects on downstream analyses.

## Introduction

Fourier Transform Mass Spectrometry (FT-MS) provides high performance in terms of sensitivity, resolution, and mass accuracy simultaneously over many other types of MS commonly used for biomolecular detection and quantification. These combined capabilities provide tangible analytical and interpretative improvements including: i) the ability to resolve isotopologues with identical unit masses but different real masses enabling multi-element isotopic natural abundance correction (H. N. Moseley, 2010) (Carreer, Flight, & Moseley, 2013) in multiple labeling experiments (Yang, Fan, Lane, & Higashi, 2017); ii) improved assignment accuracy (although often not fully unambiguous assignments) (Kind & Fiehn, 2006); and iii) the detection of low concentration metabolites in the subfemtomolar range (Eyles & Kaltashov, 2004) (Dettmer, Aronov, & Hammock, 2007). In the metabolomics field, these improvements permit more complicated, but more informative experimental designs such as the use of multiple isotope-labeled precursors for use in stable isotope-resolved metabolomics (SIRM) experiments (Yang et al., 2017). These experiments can provide the information needed to elucidate unknown metabolic pathways (Creek et al., 2012) (Higashi, Fan, Lorkiewicz, Moseley, & Lane, 2014), quantify the relative flux through metabolic pathways (Hiller, Metallo, Kelleher, & Stephanopoulos, 2010), identify multiple metabolite pools (T. W. M. Fan et al., 2012), and identify active metabolic pathways under various cellular conditions (Sellers et al., 2015). In turn, these informational gains enable researchers to build more complete models of cellular metabolism and to better understand life processes, both healthy and pathological, more completely at a mechanistic level. These models facilitate the identification of potential targets for pharmaceutical intervention (T. W. Fan et al., 2009) and the quantification of differential drug response at a molecular level (Harris et al., 2012).

While the potential benefits of FT-MS are significant, so are the limitations. In addition to the increased cost and complexity of the instruments, the amount of data produced by a single run on an FT-MS instrument can be quite large and when deployed in a high-throughput environment as with all high data capture ‘omics techniques, necessitates the use of automated tools for data reduction, data quality assessment and control, peak (feature) assignment, and other downstream analyses enabling information extraction and interpretation within reasonable timeframes. In particular, the extraction and interpretation of meaningful data from MS spectra often necessitates correct peak assignment. While peak assignment is relatively straight-forward and well-validated in targeted MS experiments, especially when combined with chromatography and/or done in tandem (Ogura, Bamba, & Fukusaki, 2013) (Zhang & Brodbelt, 2004) (Astarita, Ahmed, & Piomelli, 2009), assignment of untargeted MS analyses of non-polymeric biomolecules remains harder to perform in a rigorous, error-limiting manner, even with the advanced capabilities of FT-MS.

In untargeted FT-MS metabolomics approaches, m/z database-based assignment tools such as LipidSearch (Peake, Yokoi, Wang, & Yingying, 2013) and PREMISE (in-house tool developed within the Center for Environmental and Systems Biochemistry (CESB)) have been used to assign FT-MS features observed in direct infusion experiments (Lorkiewicz, Higashi, Lane, & Fan, 2012) (Yang et al., 2017). These database approaches have the advantage of low computational overhead and *a priori* knowledge when the m/z database is tailored to the biological system being studied. However, tailored m/z databases limit discovery – a stated goal of many untargeted analyses (H. N. B. Moseley, 2013), introduce assignment bias, and have difficulty disambiguating possible assignments. Also, these assignment approaches are often based on the matching of a single spectral feature to an m/z value in the database, which are statistically error-prone due to a lack of aggregate, cross-validating evidence.

However, the ability to identify true signal is essential for the accurate assignment and interpretation of all MS datasets, including FT-MS datasets, as all analytical techniques have the potential to generate artifactual signals. Artifactual signals are signals not representative of the sample composition but rather arise from instrumental or data processing limitations. Such signals obviously do not represent the underlying biochemistry of a sample, and at best complicate data interpretation and at worst lead to incorrect interpretations. As the scale of experiments increase and untargeted analyses become increasingly prevalent, as is the case with the field of metabolomics (Goodacre, Vaidyanathan, Dunn, Harrigan, & Kell, 2004), the ability to distinguish sample-related signal and artifactual signal becomes increasingly important.

Fundamentally, artifactual MS peaks can be divided into two major types. First are artifactual peaks that result from unexpected ions during spectral acquisition. For example, the presence of contaminant compounds (Keller, Sui, Young, & Whittal, 2008) (e.g. plasticizers and keratin) and spontaneous chemical reactions during analysis (e.g. molecular rearrangements (McLafferty, 1959)) are just two of many mechanisms by which unexpected ions can be produced and subsequently detected by a spectrometer. In this case, the peak signals represent real analytes; however, these detected analytes are not representative of the sample. The second type of artifactual peaks are those that do not correspond to ions present during spectral acquisition. The cause of these second types of artifacts typically depends upon the mass spectrometry platform on which the spectrum was acquired. Examples for FT-MS will be described below. Also, characteristics of real peaks can be artifactual. For example, ion suppression and overloading an ion trap can distort acquired spectral intensities.

The use of the Fourier transformation in the processing of FT-MS raw data introduces new avenues for generating artifactual peaks in resulting FT-MS spectra. This is generally well known, especially for other analytical techniques that utilize the Fourier transform with common FT-specific artifacts like sinc wiggles in NMR (Hore, 1985) and side lobes in FTIR (Griffiths & Pariente, 1986). However, previous studies regarding FT-MS specific artifacts have focused almost exclusively on “harmonic peaks” (Mathur & O’Connor, 2009) and “sidebands” or “peak ringing” (Miladinović, Kozhinov, Tsybin, & Tsybin, 2012), which do not cover all the artifact types we have observed in our FT-MS spectra. We have observed two additional FT-MS artifacts which we have named fuzzy sites and partial peak ringing.

In summary we have observed three distinct artifact types in our FT-MS spectra that have the potential to confound meaningful assignment and interpretation of FT-MS datasets. Furthermore, of the three observed artifact types only one appears to have been previously mentioned in literature. We have developed tools for detecting these artifacts and mitigating their effects on data interpretation.

## Materials and Methods

### HPD Artifact detection

Although the three FT-MS artifacts outlined above (ringing, partial ringing and fuzzy sites) have distinct appearances, they all result in regions of high peak density (HPD). These artifacts can be collectively referred to as HPD artifacts due to this shared property and we leverage this property to create an automated tool for their detection. Our HPD-detector is implemented using the Python programming language (Van Rossum & Drake Jr, 1995) version 3.4 and makes extensive use of the Numpy library (Walt, Colbert, & Varoquaux, 2011) to accelerate calculations. Starting with a peaklist in a Javascript Object Notation (JSON) format (Supplemental Figure 2), the detector first parses and sorts the peaks in ascending order of their m/z values, enabling the use of algorithmically efficient methods for searching the peaklist by m/z, namely a binary search. Next, a 1 m/z window is slid across the spectrum in 0.1 m/z increments. At each increment, a pair of binary searches are used to find the peaks within the window that are then counted to give a peak density metric (the value D in Step 1 of Figure 1) that is assigned to the mean m/z of the window.

**Figure 1:**
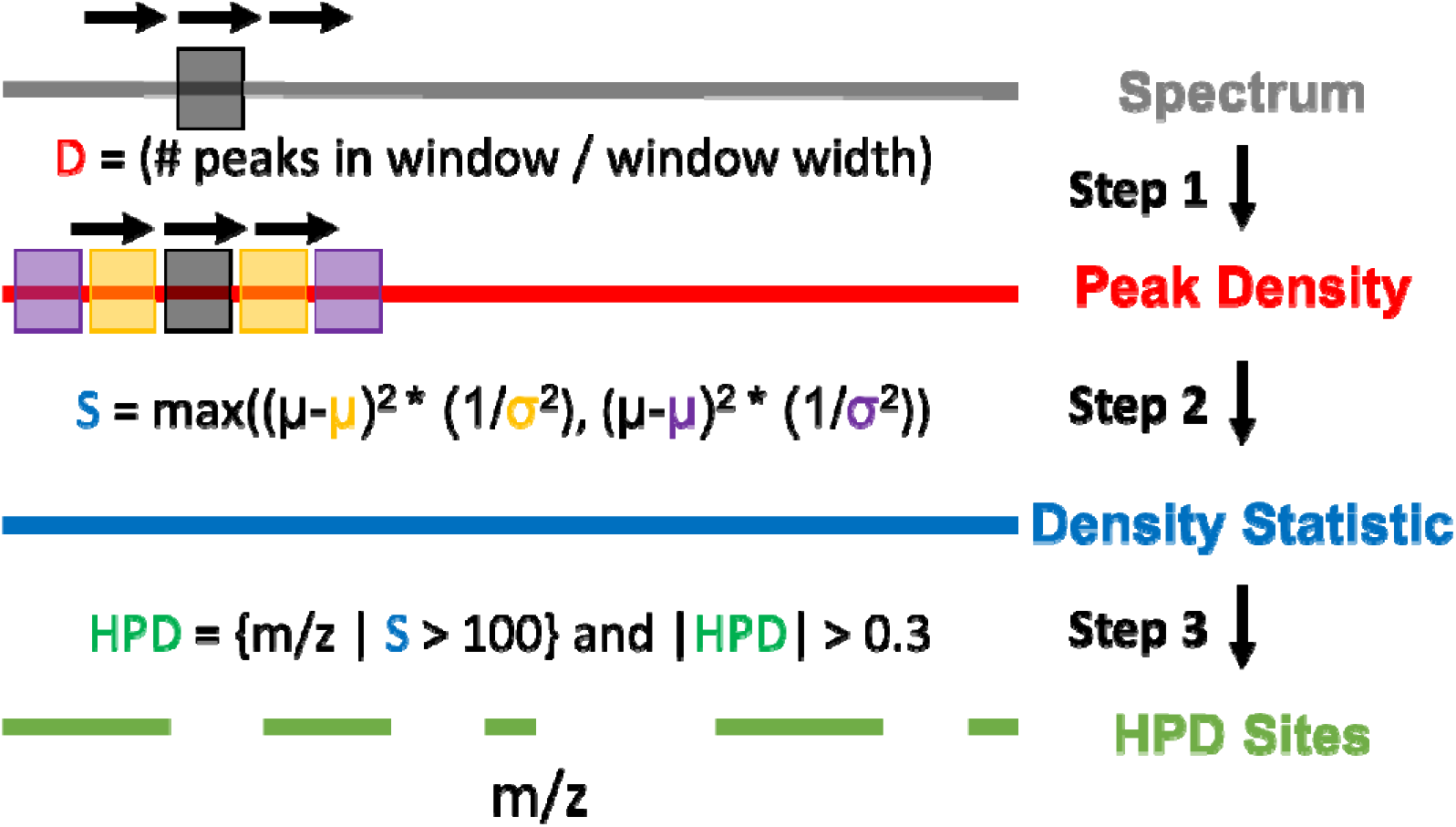
Automated HPD-Site Detection. The HPD-site detection algorithm requires three steps. First, a peak density metric is calculated for the spectrum using a sliding window method. Second, a set of K+1 windows and the peak density metric are used to calculate a density statistic for each portion of the spectrum. This metric flattens out density differences due to signal-to-noise differences or baseline differences and highlights spectra with HPD artifacts (Fig 2E, F, G, H). Filtering this metric reveals the location of the HPD artifacts. The color of variables in the equation above reflect which window or step in the algorithm from which they were derived.

By comparing these peak density statistics to one another, HPD artifacts can be found. This comparison requires applying another window operation to the density metrics calculated previously. In this operation, N pairs of non-overlapping ‘reference’ windows distributed symmetrically around a single ‘test’ window are moved across the spectrum in 0.1 m/z increments. The test window (the single black box in the peak density plot in Figure 1) is the region of spectrum being tested for HPD phenomena using the reference windows as estimates of “normal” peak densities (the yellow and purple boxes flanking the black box). At each increment, the mean and standard deviation of the peak density in the test window and each pair of reference windows are calculated. Each window is 3 m/z in width. The mean of the test window is then compared to the mean of each pair of reference windows and normalized by the standard deviation of the corresponding pair of reference windows yielding N chi-squared-inspired statistics for each test window and the max value is assigned to the mean m/z of the test window (Step 2 in Figure 1). Taking the maximum comparison value across multiple pairs of reference windows increases the sensitivity of the test and insulates the method against the edge case where all reference windows and the test window contain HPD artifacts and thus do not appear statistically different from one another. Although higher values of N are theoretically superior, testing showed no significant improvement for N > 2.

In the final step, the continuous subdomains of m/z space at least 0.3 m/z in width with density statistic values over 100 are reported (Step 3 in Figure 1). These regions likely contain some form of HPD phenomena as they have significant differences in peak density compared to neighboring regions.

### Ringing detection

The HPD-detector outlined above detects fuzzy sites and some instances of ringing and partial ringing, but has lower sensitivity for ringing artifacts. Partial ringing often does not increase peak density sufficiently to be detected by the HPD-detector, while true ringing can often occupy a very wide m/z window, making their detection more difficult. Therefore, we created a dedicated ringing detector in Python 3.4 that identifies sets of peaks that are possibly assignable to the same compound and has matching m/z differences to other expected compounds.

The ringing detector requires as input a peaklist in JSON, a list of expected m/z values in JSON (Supplemental Figure 3), and a description of the adducts to consider for each ion (Supplemental Figure 4). The provided list of expected m/z values was built using the MaConDa database (Weber, Li, Bruty, He, & Viant, 2012) of common mass spectrometry contaminants and each entry represents the non-adducted monoisotopic form of a contaminant while the adduct and isotopologue file was created manually.

In the first step of the analysis, the ringing detector expands the list of expected m/z values by generating their adducts and a set of expected isotopologues as specified by the adduct and isotopologues file. This expansion ensures that we are searching for the most likely forms of the provided compounds. With the expanded set of expected m/z values calculated, the detector parses the peaklist and sorts the peaks in ascending m/z order to enable binary searching by m/z value. Next, an estimate of the noise intensity level is calculated. From an ascending list of peak intensities, the ringing detector first identifies the first quartile of the intensity values, i.e. the lowest quartile of intensities, and copies them to a noise intensity list. While the standard deviation of the noise intensity list is less than 1.2 times the mean of the noise intensity list, the next 100 intensities are copied to the noise intensity list. Once this process is complete, the median of the noise intensities is calculated. Five times the median noise intensity is considered the intensity cutoff for the spectrum and intensities above the cutoff are very likely “true” signal. A noise sample is then generated by selecting a random set of peaks below the noise cutoff equal in size to the number of peaks above the cutoff.

In this noise sample, the ringing detector searches for the expected m/z values and for every expected m/z in the set of expanded m/z values, a binary search is used to find all peaks within +/−0.05 m/z of the expected m/z value. The m/z difference between each found peak and all other peaks in the noise sample is calculated (peak mass difference vector), as is the m/z difference between the expected m/z and all other expected m/z values (query mass difference vector). The number of matching m/z differences within a match-difference m/z tolerance between the peak mass difference vector and the query mass difference vector are counted. For the ultra-high resolution FT-MS spectra analyzed in this study, we used a tight match-difference tolerance of ±0.00005 m/z. The peak with the most matching m/z differences and the number of matches is reported. Since these are noise peaks, these matches are unlikely to be correct and the number of m/z difference matches is due to random chance. Therefore, the ringing detector uses this sample of mostly false matches to estimate a cutoff for accepting true matches. The detector calculates this cutoff as the mean of noise matching count plus three times the standard deviation of the distribution of noise matching counts.

Finally, the ringing detector repeats the search procedure using the entire peaklist. During this search, only peaks with many matching m/z differences greater than the match cutoff calculated from the noise peaks are reported as assignments. When ringing or partial ringing occurs, multiple assignments occur for an expected m/z value as there are many peaks close to the primary peak that have similar m/z differences to other expected compounds.

### FT-MS instruments

To determine the instrument dependence of various artifacts described here, spectra from several different FT-MS instruments were used. The first set of instruments were three Thermo Tribrid Fusion instruments. Fusion 1 and Fusion 2 are maintained by the Center for Environmental and Systems Biochemistry (CESB) at the University of Kentucky, while Fusion 3 (Serial # FSN 10144) is maintained by National Resource for the Mass Spectrometric Analysis of Biological Macromolecule at the Rockefeller University. Fusion 1 (Serial # FSN10115) was delivered to CESB in October 2013. In March of 2016, Fusion 1 had its firmware upgraded to the most recent version at that time. Pre-firmware upgrade Fusion 1 (Fusion 1 – Before) has different HPD artifact patterns than post-firmware Fusion 1 (Fusion 1 – After). Fusion 2 (Serial # FSN10352) was delivered to CESB in May 2015 with the upgraded firmware. Spectra from a Thermo Scientific Orbitrap Fusion Lumos Tribrid mass spectrometer (Serial # FSN20208, delivery data July 2016), which is maintained by the Proteomics Resource Center at the New York University Langone Medical Center, were also examined. The Lumos represents an improvement upon the original version of the Tribrid Fusion. In addition to the Fusion instruments, we examined spectra from a Thermo Q-Exactive+ instrument, which is another Orbitrap instrument maintained by High Resolution Metabolomics Laboratory (HRML) at the Institute of Biological, Environmental and Rural Sciences at the Aberystwyth University in the United Kingdom. Also, spectra from a Bruker Solarix instrument (Serial # 150506 A), an ICR-type FT-MS delivered to CESB in April 2014, were examined.

### Samples analyzed by FT-MS

#### Sample A: Solvent Blanks with and without Avanti Lipid Standards

The solvent blank was composed of Isopropanol:MeOH:Chloroform 800μl:344μl:200μl. The solvent blank was mixed with 28μl 1 M ammonium formate (final ~20 mM; Aldrich #516961), and without or with 70μl Avanti SPLASH™ Lipidomix^®^ Mass Spec Standard (cat# 330707) in MeOH. The solvent blank without or with lipid standards was loaded onto a 96-well polypropylene PCR plate (USA Scientific cat# 1402-9800) and 15 μl was injected into the Fusion 1 by direct infusion through an Advion nanomate. Various resolution and microscan settings were tested for the Orbitrap mass analyzer in positive mode with 7 min acquisition, normal mass range between 150-1600 m/z, S-lens RF level 60%, AGC target 1e5, maximum injection time 100 ms, and Easy-IC on.

#### Sample B: Mouse Liver ICMS Standard

Mouse livers were excised from NSG mice within 5 minutes of euthanization, and flash frozen in liquid nitrogen and ground into powder under liq. N_2_ to <10 um particles. Approximately 0.5 g of the powder was extracted with 50 mL Acetonitrile: water (6:4, v/v). After centrifugation at 22 k rpm and 4 °C for 20 min, the supernatant containing polar extracts was distributed into aliquots and lyophilized for long-term storage at −80 °C. Immediately before injection, the lyophilized powder was reconstituted with water and 10 μL was injected onto an ICS5000+ system (Dionex) interfaced to the FT-MS (Fusion 2). Data were acquired in negative mode at a resolving power of 500,000 (at m/z=200) over 52 min of chromatography. The mass range was set between m/z 80-700, maximum injection time was 100 ms with 1 microscan, AGC target was 2e5, S-lens RF level was 60%, and Easy-IC was turned on for internal mass calibration.

#### Sample C: ECF Solvent Standard

The ECF solvent blank was composed of acetonitrile: water 9:1 (v/v) with a concentration of 20 μM NaCl to convert positively charged ions into sodium adducts (Yang et al., 2017).

#### Sample D: Paired Human NSCLC Cancer and Non-Cancer Tissue Samples

The samples analyzed are the lipids extracted from flash frozen resected lung tissues from human subjects with resectable stage I or IIa NSCLC (non-small cell lung cancer) collected under a University of Louisville approved Internal Review Board (IRB) protocol. Paired cancerous and non-cancerous tissue 5 cm form the tumor margins in the same lobe of the lung were cut by the surgeon and immediately flash frozen in liq. N_2_, and stored at <-80°C prior to metabolite extraction as previously described (Sellers et al., 2015). Samples were pulverized under liq. N_2_ to <10 um, and extracted using a modified Folch method as previously described (Ren et al., 2014). 1 mM butylated hydroxytolune was added to the lipid layer and then dried by vacuum cetrifugatin at room temperature. Samples for FT-MS analysis were redissolved in isopropanol/methanol/chloroform 4/2/1 (v/v/v) with 20 mM ammonium formate (95 μl of solvent, 5 μl of sample).

#### Mass Spectrometry Analysis of Samples C and D

Ultrahigh resolution (UHR) mass spectrometry was carried out on a Thermo Orbitrap Fusion interfaced to an Advion Nanomate nanoelectrospray source using the Advion “type A” chip, also from Advion, inc. (chip p/n HD_A_384). The nanospray conditions on the Advion Nanomate were as follows: sample volume in wells in 96 well plate – 50 µl, sample volume taken up by tip for analysis – 15 µl, delivery time – 16 minutes, gas pressure – 0.4 psi, voltage applied – 1.5 kV, polarity – positive, pre-piercing depth – 10 mm. The Orbitrap Fusion Mass Spectrometer method duration was 15 minutes, and the conditions during the first 7 minutes were as follows: scan type – MS, detector type – Orbitrap, resolution – 450,000, lock mass with internal calibrant turned on, scan range (m/z) – 150 – 1600, S-Lens RF Level (%) – 60, AGC Target – 1.0e^5^, maximum injection time (ms) – 100, microscans – 10, data type – profile, polarity - positive. For the next 8 minutes, the conditions were as follows for the MS/MS analysis: MS properties: detector type – Orbitrap, resolution – 120,000, scan range (m/z) – 150 – 1600, AGC Target – 2.0e^5^, maximum injection time (ms) – 100, microscans – 2, data type – profile, polarity – negative; monoisotopic precursor selection – applied, top 500 most intense peaks evaluated with minimum intensity of 5e^3^ counts; data dependent MS^n^ scan properties: MS^n^ level – 2, isolation mode – quadrupole, isolation window (m/z) – 1, activation type – HCD, HCD collision energy (%) – 25, collision gas – Nitrogen, detector – Orbitrap, scan range mode – auto m/z normal, Orbitrap resolution – 120,000, first mass (m/z) – 120, maximum injection time (ms)-500, AGC target – 5e^4^, data type – profile, polarity – positive. The ion transfer tube temperature was 275 °C. (Yang et al., 2017).

#### Sample E: Human Plasma

Pooled lithium heparin treated plasma (Seralab) was extracted using the methods described by Koulman et al, 2017 (Acharjee et al., 2017). Briefly, 15 µL of plasma was extracted with 100 µL of ultra-pure H2O in a glass vial (2 mL). 250 µL of MeOH was added, and lipids were partitioned into 500 µL of Methyl-tertiary-butyl ether. Following centrifugation (13,000 rpm, 4°C, 4mins), a 20 µL aliquot of the organic layer was then transferred to a 96-well glass coated plate (ThermoFisher). 95 µL of a solution containing 7.5mM ammonium acetate in IPA:MeoH [2:1] was also added to the well. Direct infusion high-resolution mass spectrometry was performed using on a Q-Exactive+ Orbitrap (Thermo), equipped with a Triversa Nanomate (Advion). The Nanomate infusion mandrel was used to pierce the seal of each well before analysis, after which, with a fresh tip, 5 μL of sample was aspirated, followed by an air gap (1.5 μL).

## Results

### Manual Investigation of Artifacts

Before developing automated tools for their detection, many spectra were manually inspected and artifacts with HPD properties (Figure 2) were observed and characterized (Figure 3). Figure 3 illustrates three major types of HPD artifacts observed: fuzzy sites, peak ringing and peak partial ringing and all three appear distinct at the aggregate and scan level. All three artifacts are obvious upon manual inspection, but the growing popularity of FT-MS-based experiments necessitates automated methods for their efficient identification.

**Figure 2:**
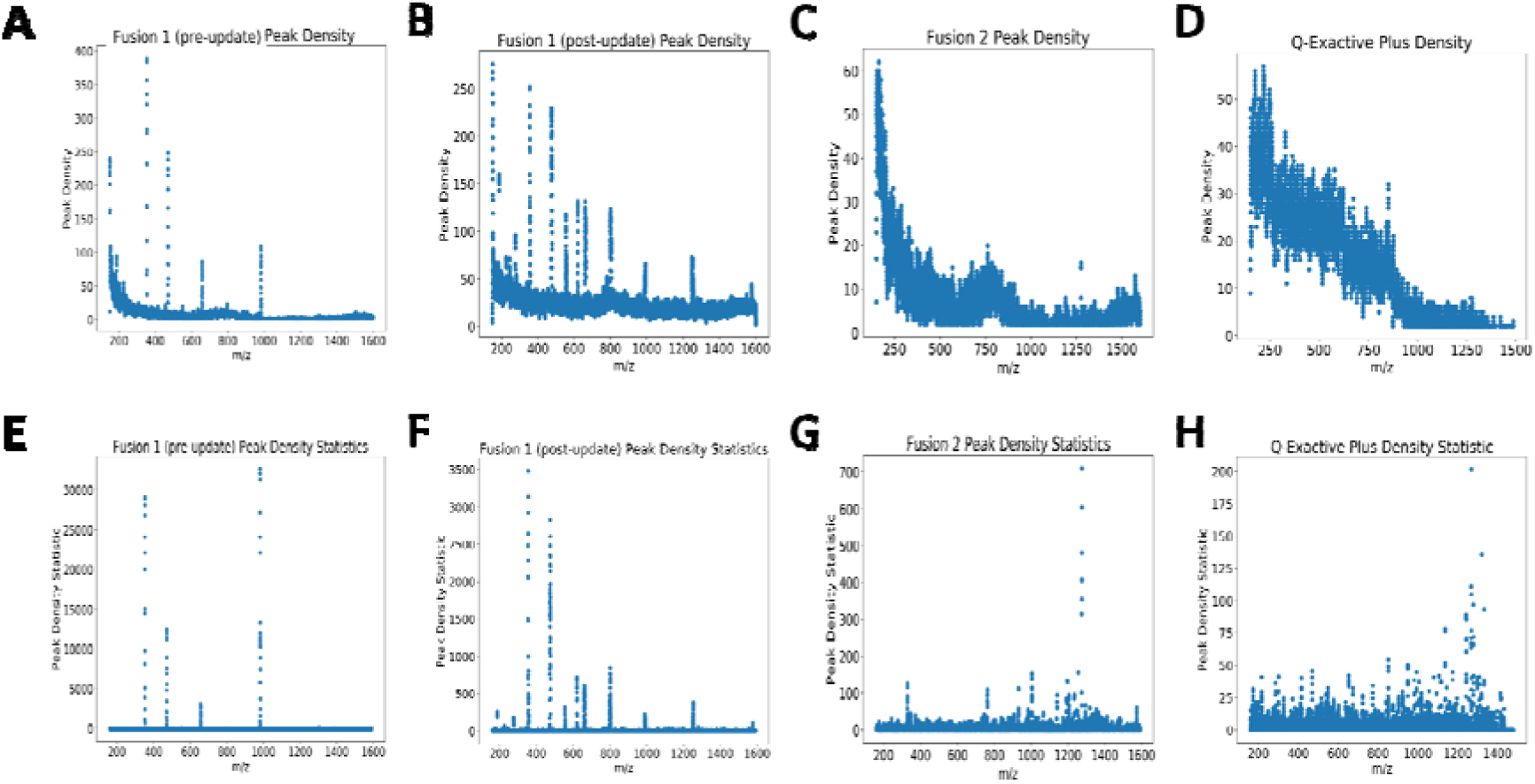
Peak Density and Peak Density Statistics. Peak density metric plots produced by our HPD-detector tool highlight the impact of the instrument on peak density and HPD artifact location. All instruments have higher peak densities at lower m/z representing trends in signal-to-noise with respect to m/z in FT-MS. The sharp spikes in peak density correspond to HPD artifacts. The location of these spikes on Fusion 1 is different before and after the firmware update (A, B), suggesting instrument-level processing of the data is related to HPD generation. E-H show the effectiveness of our peak density statistic metric for flattening the non-constant baseline observed in plots of the raw peak density. Without this correction, identifying HPD regions reliably is difficult. The presence of severe HPD regions in effectively all of our tested Fusion spectra contrasts with the lack of severe HPD results in Q Exactive spectra supports an instrument level explanation for HPD phenomena. A, B, C, E, F, G were generated from spectra acquired using sample C. D and H were acquired using sample E.

**Figure 3:**
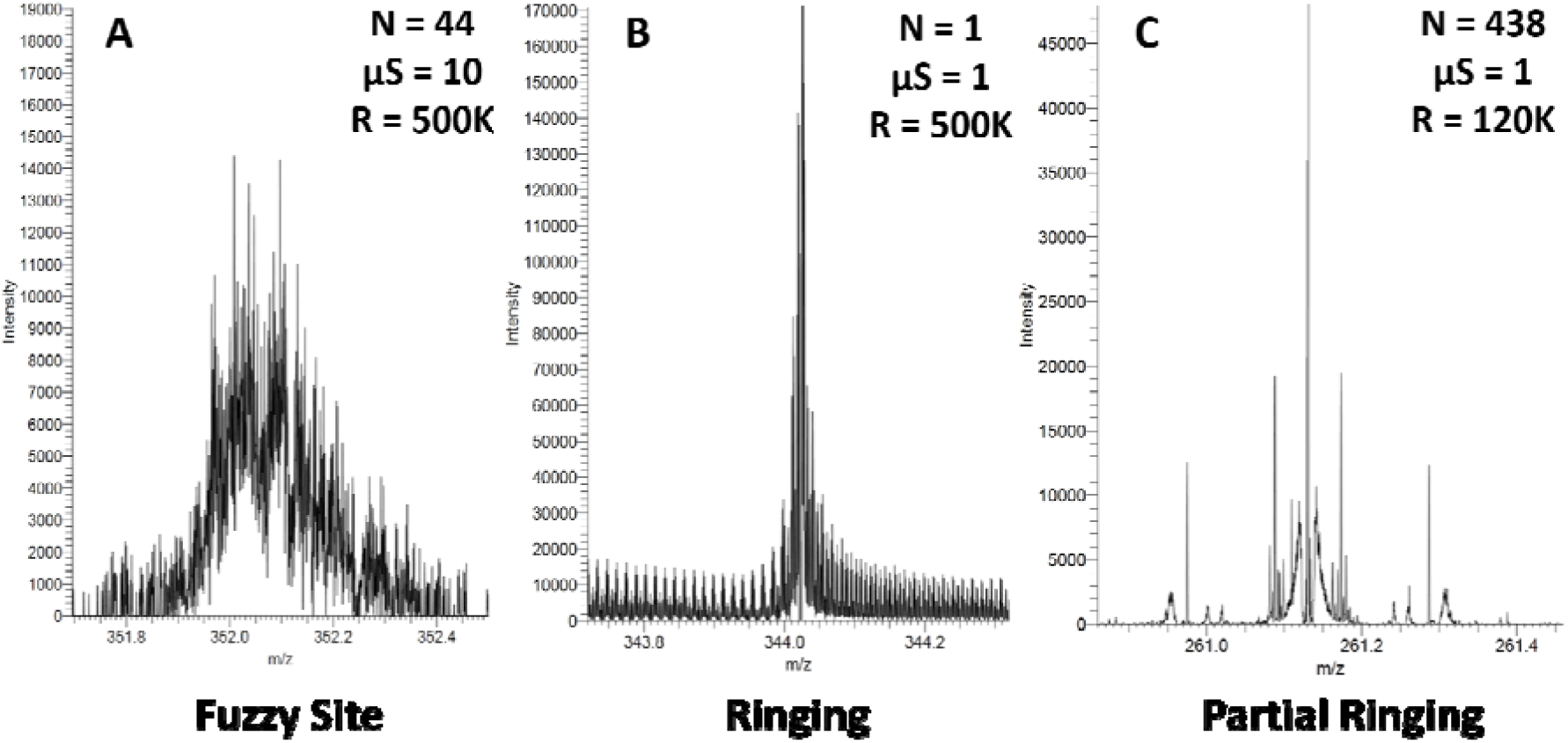
Three Types of HPD Artifacts. There are three subclasses of HPD artifacts we have investigated. The first is the fuzzy site which we believe is a novel artifact type (A, Sample A). A fuzzy site appears pseudo-Gaussian at the aggregate level. Second is ringing, a well-known FT-MS artifact where a single intense peak has many side peaks (B, Sample B). We only observed ringing at the scan level. The third artifact is partial ringing which is a ringing-like artifact at the aggregate level (C, Sample C). Partial ringing appears like ringing but has not been discussed in the literature.

### General HPD detection across FT-MS instruments

Using the HPD artifact detector, we generated plots of peak density for a variety of example spectra across various FT-MS instruments (Figure 2). For most Orbitrap instruments, the peak density decreases monotonically with respect to m/z. While the same general decreasing trend is observed in our ICR instrument and a Lumos Orbitrap instrument, the overall curve is not monotonic (Supplemental Figure 1). These trends are partially explained by differences in signal-to-noise ratio at different m/z values and between different instruments. The peak density properties demonstrate that a statistical approach is necessary to identify HPD as there exists no sufficiently sensitive and selective cutoff on raw peak density for all m/z or all instruments. However, our statistical approach effectively normalizes the base peak density in the regions of spectra we are testing and, in turn, compensates for changes in baseline peak densities revealing regions of significantly higher peak densities (Figure 2E-H).

Although fluctuations in peak densities are expected due to differences in the distribution of compounds in m/z space, this fails to explain the massive difference in peak density statistics present in the spectra and similar patterns are observed in blanks and in scans from failed injections, which resulted in little analyte or solvent. Furthermore, analytical samples derived from the same biological sample have HPD regions at different locations with different instruments and the location of these artifacts differs before and after a firmware update on the Fusion 1 machine (Figures 2A, 2B, 2E, 2F). Manual inspection of a subset of the detected HPD regions consistently failed to identify patterns between peaks that are explainable by chemical phenomena (*e.g.* isotopologues, different charges, etc.). Together, these findings support an artifactual basis for these regions of spectra and suggest an instrument-level effect leading to their production.

### Detection and characterization of fuzzy sites

The fuzzy site artifacts were first observed and described as “ugly sites”. At the aggregate spectrum level (Figure 3A, 4A,C), fuzzy sites have HPD characteristics and a pseudo-Gaussian distribution of peak intensities between the noise baseline and presumed signal peaks. The intermediate intensities of these peaks make identifying and filtering these regions by intensity alone difficult. Fuzzy regions, like other HPD artifacts, have peak m/z differences that are not explainable by isotopologue, charge, or harmonic patterns. A typical fuzzy site occupies a small m/z window (e.g., 0.5 to 1.5 m/z approximately), with larger ranges typically occurring at higher m/z values. Fuzzy sites rarely occur singly: many fuzzy sites are typically observed in a single spectrum. Collectively, these sites can represent a significant portion of the peaks over a much smaller amount of the total m/z range. Fuzzy site location varies between analytical replicates on the same instrument and with sample composition (Supplemental Figure 6). Fuzzy sites have been observed in samples with failing or no injection as well.

Fuzzy sites also have interesting properties at the scan-level as well. While the timing between scans and injection as well as inconsistencies between injections can result in non-perfect scan-to-scan correspondence between peaks (e.g. a peak is present in scan X, but not in scan Y), peaks of the same chemical origin should appear consistently between scans near their true m/z, roughly within the resolution of the instrument. At the scan level, the peaks in a mass range identified as a fuzzy site at the aggregate level have very low peak correspondence (Figure 4B). In any given scan, only sections of the fuzzy site region will have peaks and those sections that are populated with peaks change from scan to scan. However, as increasingly more scans are averaged together, the Gaussian-like distribution of a fuzzy site at the aggregate level becomes clearer (Figure 4A,C). Fuzzy sites appear distinct from either peak ringing or partial peak ringing and represent a novel class of artifact not previously described in the FT-MS literature.

**Figure 4:**
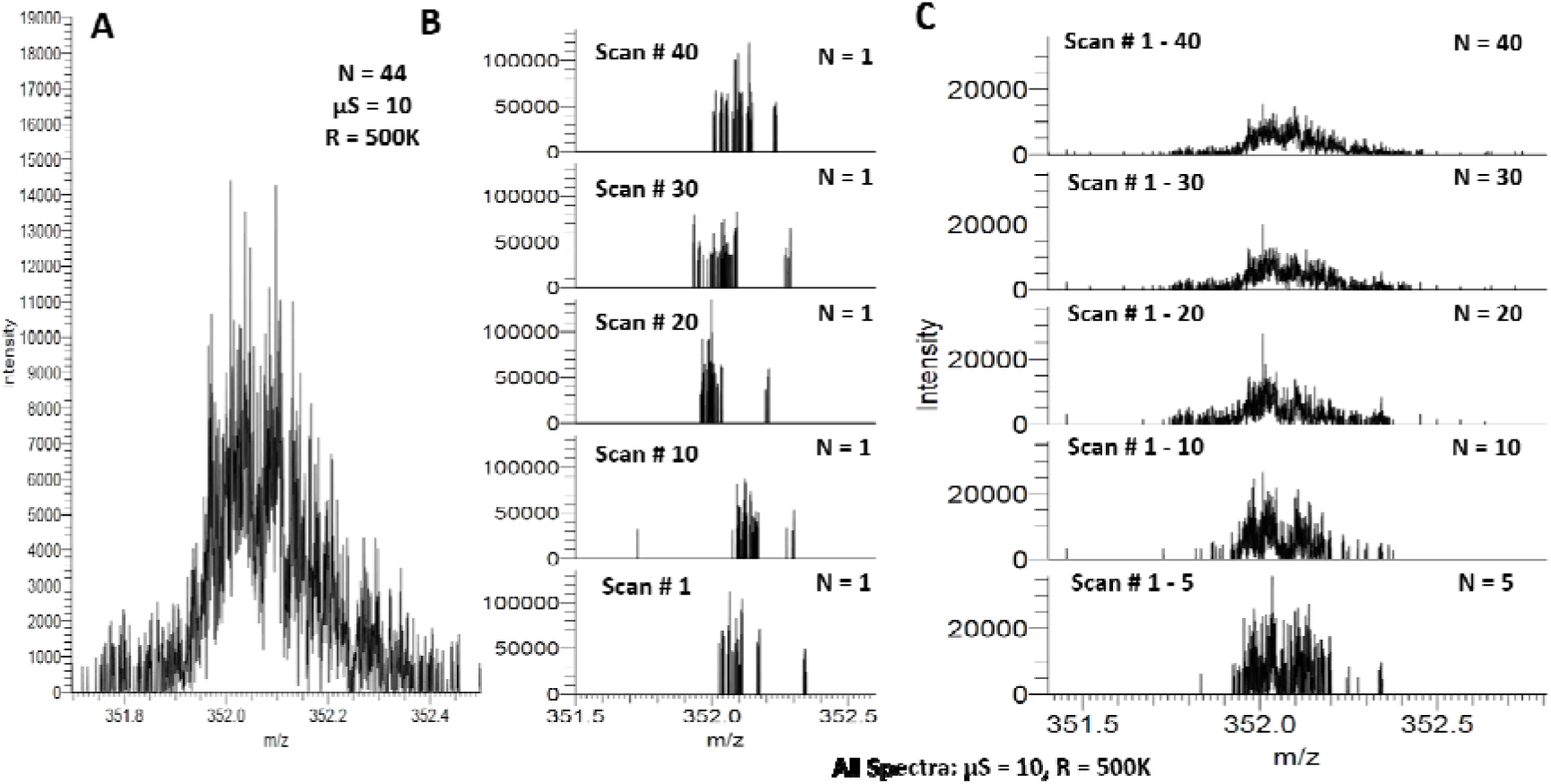
Fuzzy sites at the Aggregate and Scan Level. A typical fuzzy site (A) occupies .5 to 2 m/z at the aggregate level and has a distinct ‘fuzzy’ appearance due to very high peak density (this image is identical to 3A). At the scan level, only a subdomain of the m/z occupied by the fuzzy site contains peaks; the subdomain that is occupied varies from scan-to-scan (B). As increasingly more scans are aggregated together, the peak distribution converges to the pattern observed at the aggregate level (C). All panels were generated using Sample A.

The initial studies into fuzzy site HPD artifacts were performed on our Fusion 1 instrument where they were first observed. After developing our tools on this data, spectra from other instruments were examined for fuzzy sites to determine if these artifacts were limited to only one instrument. Spectra from the Fusion 2 instrument also contained fuzzy sites as did spectra from Fusion 3. To date, we have observed fuzzy sites in spectra from every non-Lumos Tribrid Fusion instrument examined. However, we did not find fuzzy sites in spectra examined from any other type of FT-MS instrument (Lumos, Q Exactive+, SolariX)

### Fuzzy site characteristics varies with resolution and microscan settings

With fuzzy sites found in non-Lumos Tribrid Fusion spectra, we began investigating the effect of two instrument parameters on the appearance of these artifacts. The first parameter examined was resolution, which was a parameter that was improved by the firmware update that changed the HPD properties of Fusion 1. The most up-to-date Fusion has a maximum resolution of 500K at 200 m/z (450K before update) and the second parameter was the number of microscans per scan. The number of microscans is the number of FIDs acquired and summed to create the FID that is transformed to produce the scan-level spectrum. Due to the high scan-level variability of fuzzy sites, these settings were of interest as it directly impacts how scan-level FIDs are acquired and processed.

Using a series of analytical replicates created using the same sample of solvent blank with lipid standards, spectra were acquired on the Fusion 1 instrument at 3 resolutions (120 k, 240 k and 500 k at m/z=200) and 4 microscan settings (1,2,5, and 10 microscans, but only 1,5, and 10 microscans are shown in Figure 5). No combination of settings eliminated the fuzzy sites, but they do change the general appearance of the fuzzy sites. Higher resolution results increased the peak density of fuzzy sites, indicating that the peaks within these regions may be sharper (smaller peak widths) than what is indicated at the highest resolution. Higher microscan settings increased the variability in peak intensities. Also, extremely high microscan settings (e.g. µS **=** 350) resulted in broad uniform regions for these fuzzy sites (Supplemental Figure 5).

**Figure 5:**
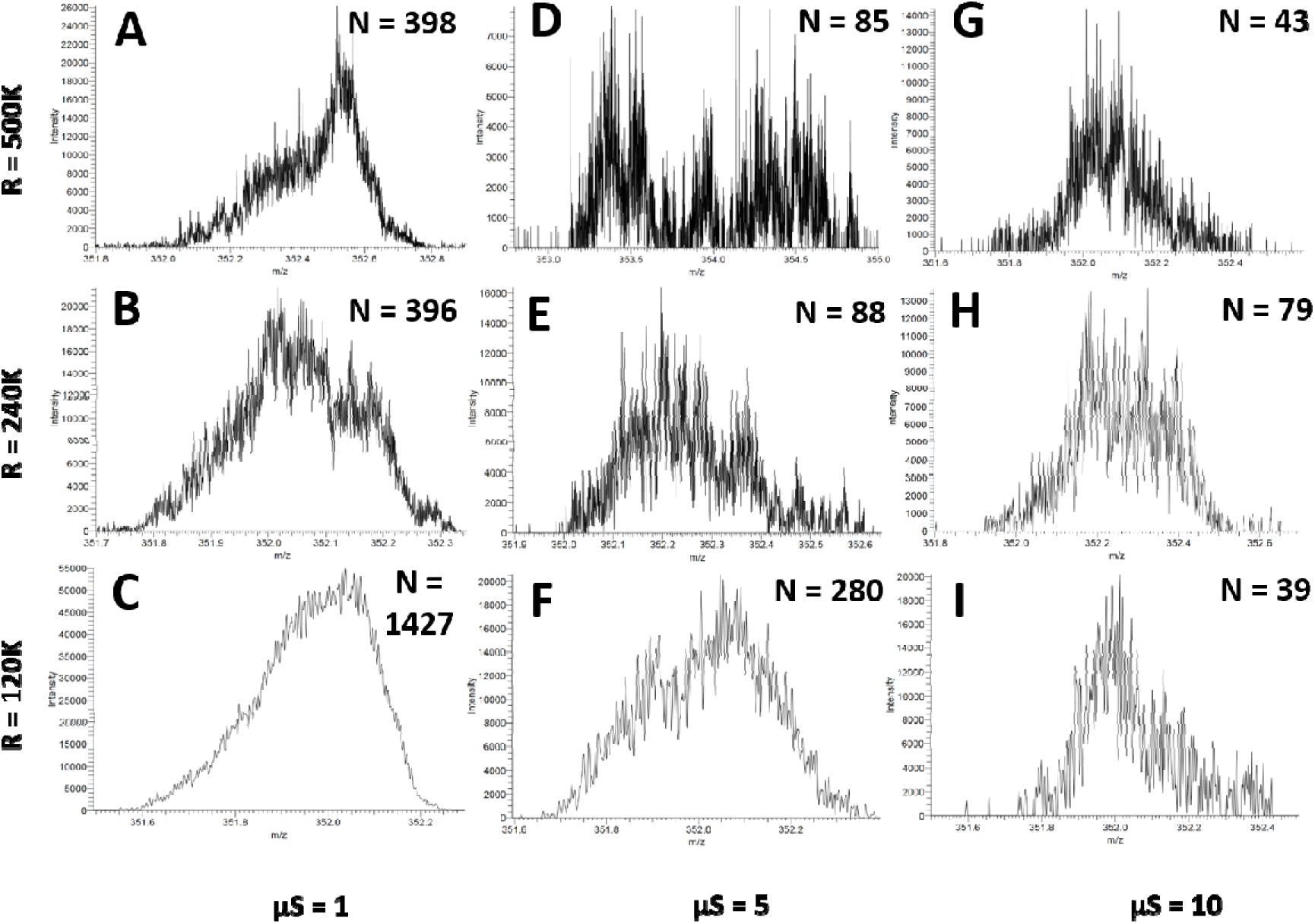
Effect of Resolution and Microscan (µS) on Fuzzy Sites. Permuting over multiple resolution and µS settings shows that no combination of tested settings eliminated fuzzy sites, but these settings do change their appearance (A-I). Number of scans collected were set so that total acquisition time was constant. Higher µS increases intensity variance with minimal impact on peak density. Increasing resolution increases peak density but has a lesser impact on peak intensity variance. All panels were generated using Sample A.

### Fuzzy site locations are sample specific, confounding these artifacts with sample class

As shown in Supplemental Figure 6, fuzzy site location varies between spectra and appears to shift significantly (a shift greater than the resolution of the instrument) with changes in sample composition. A potential hypothesis explaining this observation is that sample composition is related to fuzzy site location.

If this hypothesis is correct, fuzzy site artifacts are a potential problem for real biological applications of FT-MS. To illustrate this potential problem, consider sample D, the paired cancer and non-cancer lung tissue slices. Due to the differences in the concentrations of various metabolites between cancer and non-cancer, we would anticipate that features assigned to spectra of different sample classes could be used to distinguish sample classes. However, if fuzzy sites vary with sample class as well, artefactual features will also distinguish sample class without reflecting the underlying biochemical differences between the classes directly.

This creates a potential confounding factor in all downstream statistical analyses and reduces robustness of acquired spectra if spurious changes in sample conditions introduce new sample-specific artifacts. For example, machine learning methods such as random forest (Breiman, 2001) are trained with known classes of samples in order to classify unknown samples into one or more known classes based on spectral features identified in the training set of samples. However, these techniques rely upon an important assumption that detectable artifacts are not confounded with sample class, which may not be true. The large number of sample-specific artifactual peaks produced by these fuzzy sites can hijack the classifier training, anchoring the classification to artifacts that may change due to unforeseen sample conditions in unknown samples or changes in the analytical instrument. This effect is demonstrated in Table 1 and can greatly reduce the robustness of the classification of other samples analyzed (Table 1A). In addition, it is non-trivial to remove HPD features correctly. If a feature within an HPD region in any spectrum is just removed, this actually encodes the HPD region due to the absence of a peak (Table 1B). Therefore, the proper action is to remove the HPD-tainted feature from all spectra (Table 1C).

**Table 1:**
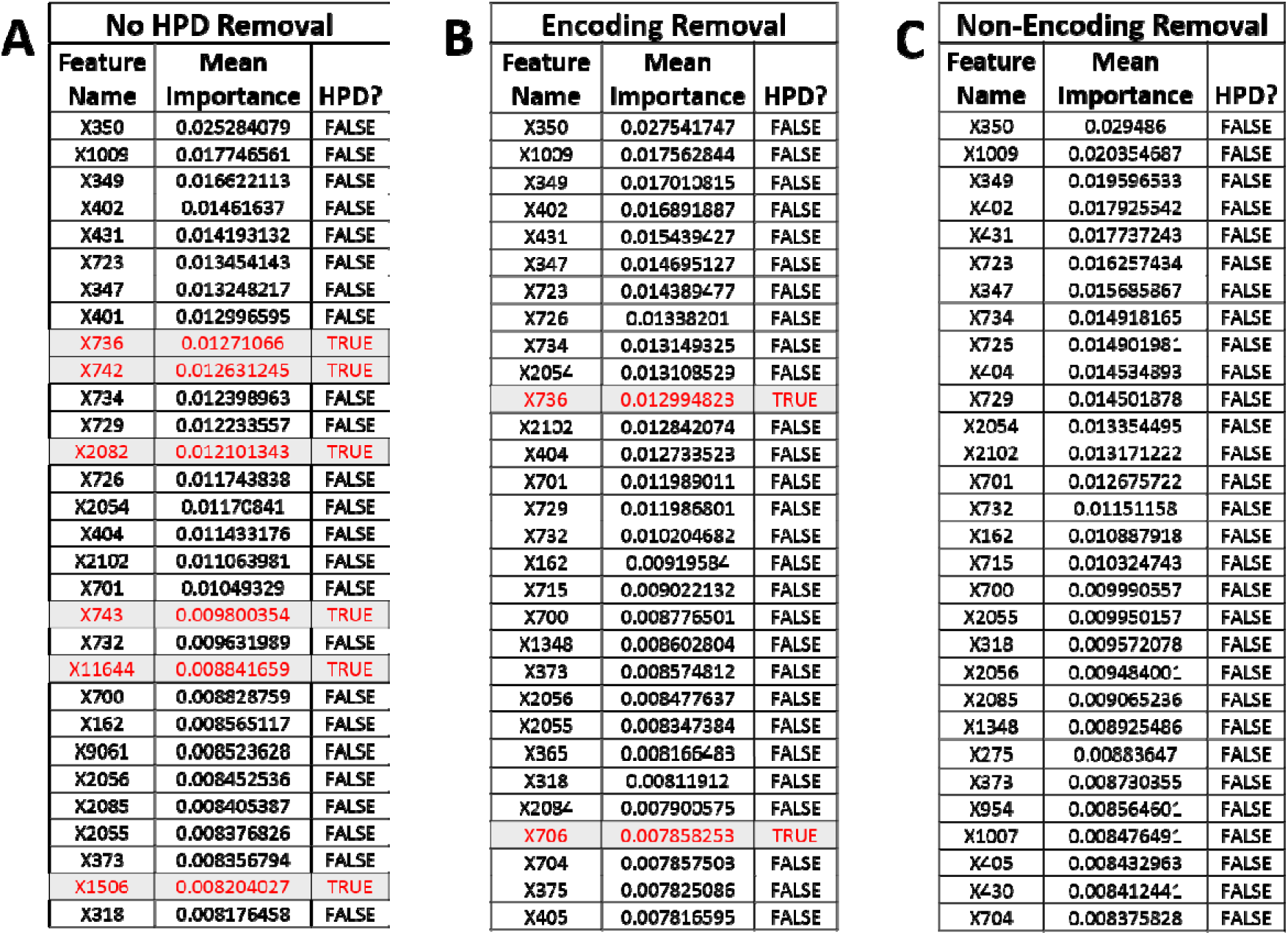
**Fuzzy Sites are Class Specific.** 20 random forests were trained using lipid search assignments from sample D and the mean importance of each feature was reported. HPD features in red were within an HPD region in at least one sample. (A) Without HPD feature removal, on average 6 out of the top 30 features are HPD features. (B) Simple removal of HPD features only within the specific HPD region encodes the HPD region into the features, which falsely boosts its importance. (C) The proper action is to remove that feature occurring within HPD regions across any spectra from all peaklists. All three classifiers could disambiguate cancer and non-cancer with 100% accuracy.

### Peak ringing

Peak ringing is an HPD artifact characterized by the presence of many peaks symmetrically centered around a very intense primary peak. The intensity of these side peaks decreases with increasing distance from the primary peak and the m/z interval between each side peak is consistent across the entire artifact. This pattern is clear in Figures 3B and 6A. Unlike fuzzy sites, where artifactual peaks appear and disappear from scan-to-scan, peak ringing is an all-or-none phenomenon at the scan level (Figure 6A); however, ringing peaks may not ring in all scans (Figure 6B) and may exhibit other artifact types (Figure 6C, D). In general, only peaks with high relative intensity in a scan will exhibit ringing behavior. Peak ringing is a well-known artifact type in FT-based instruments and can be caused by Fourier transformation of a truncated FID (Wood & Mark Henkelman, 1985). We hypothesize that this is the cause in our examples as well. There are well-known solutions to suppressing these artefacts using processing in the time domain (Guan & Marshall, 1997).

**Figure 6:**
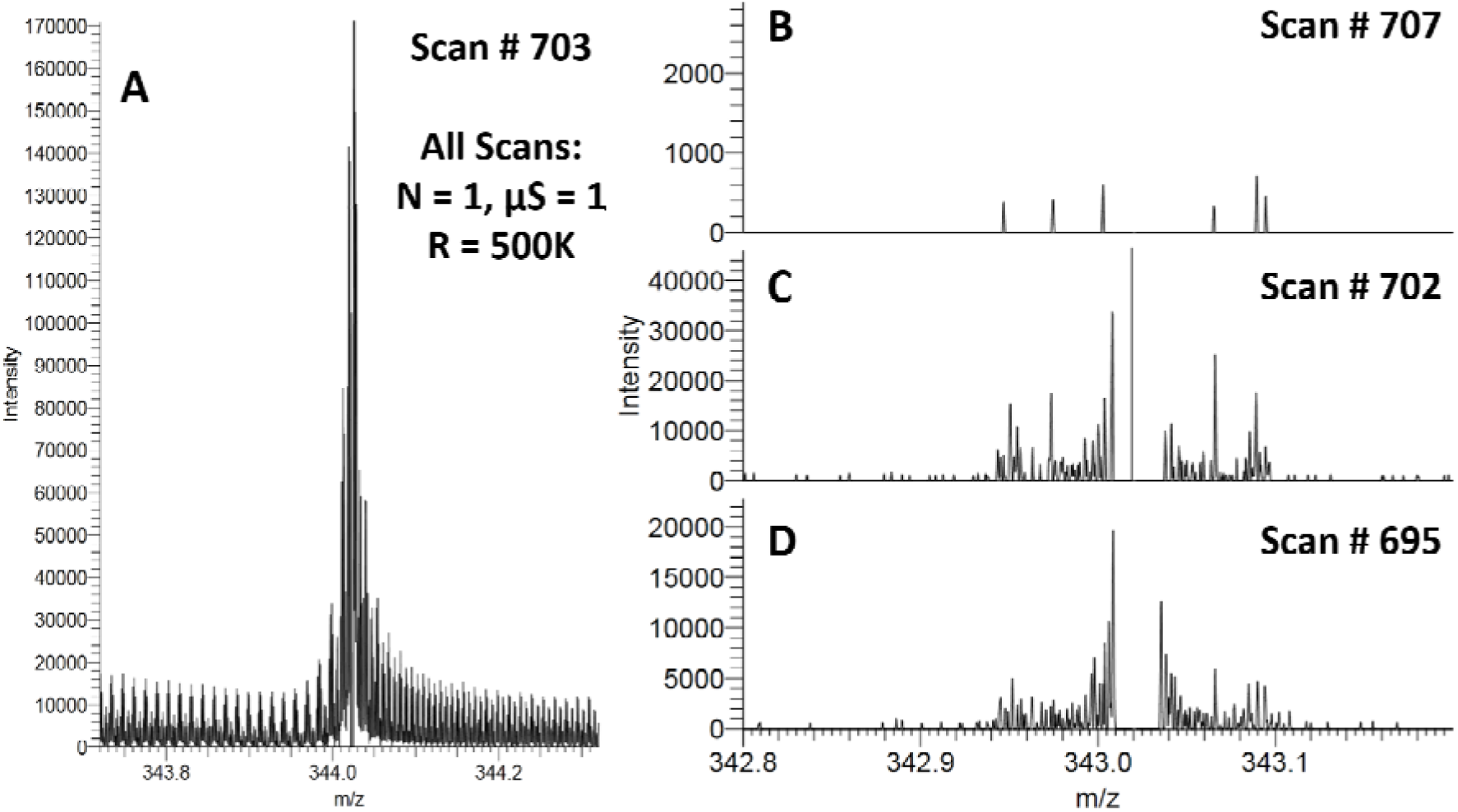
Ringing Artifacts. Ringing results in many artifactual peaks for an intense peak at the scan-level with a distinctive pattern of decaying intensity with increasing distance from the central peak (A). A complete complement of ringing peaks is present for a primary peak in a scan with true ringing. A peak that rings in one scan, does not necessarily ring in other scans; some scans demonstrate partial ringing (C, D) while other scans show no ringing (B). All panels were generated using Sample B.

### Peak partial ringing

Like ringing, partial ringing occurs around intense peaks (*i.e.*, primary peaks) (Figure 7A) but much like fuzzy sites, the artifactual peaks vary significantly at the scan-level. The artifactual peaks appear symmetrically centered around the primary peak (Figure 7A), occupying several tenths of an m/z, but they rarely occur in the immediate vicinity of the primary peak (Figure 7B, C). Side peaks from partial ringing are of lower intensity than the primary peak, but their intensities do not necessarily decrease with increasing distance from the primary peak. Although the symmetry of the peaks and the intensity pattern is less apparent in some scans (Figure 7D), when sufficient scans are aggregated, partial ringing appears smoother and more akin to true ringing except near the main peak (Figure 7A). We speculate that partial ringing is the result of incomplete ringing suppression at the scan level.

**Figure 7:**
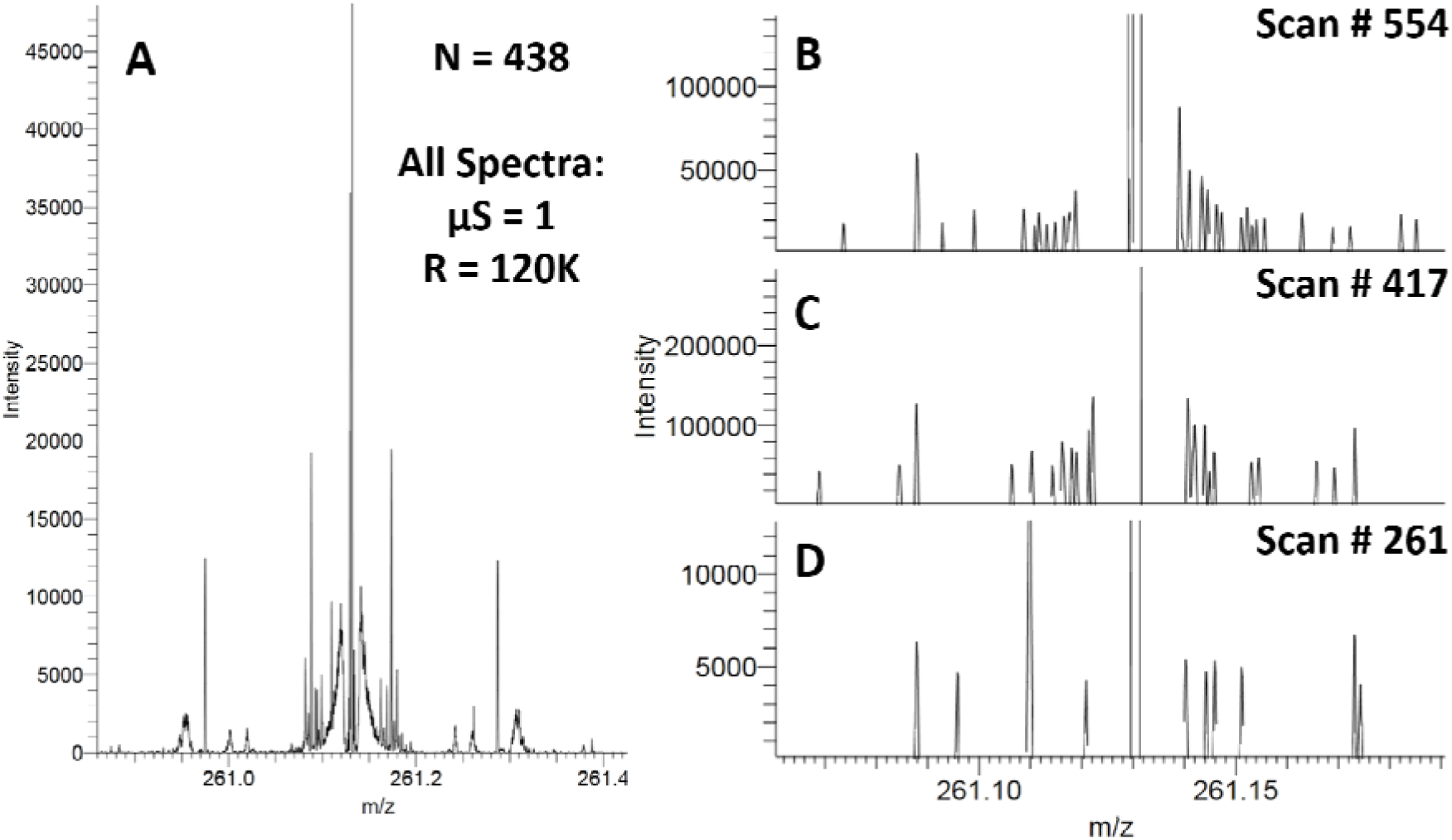
Partial Ringing. (A) Partial ringing produces peak patterns at the aggregate-level that are similar to ringing at the scan-level. Unlike ringing, partial ringing does not strictly decrease with increasing distance from the main peak and partial ringing is often absent (or greatly diminished) near the main peak. (B,C,D) At the scan level, the location of the artifactual peaks is highly variable. All panels were generated using Sample C.

## Discussion

### Origin of FT-MS artifacts

FT-related artifacts are ubiquitous in essentially all analytical techniques that leverage FT to process data. A common cause of ringing artifacts (Fig. 3B & 6) is the transformation of a truncated FID and/or improper apodization and these artifacts have analogs in both FT-NMR (*i.e.* sinc-wiggles (Hore, 1985)) and FT-IR (*i.e.* sidelobes (Philip B Tooke, 1989) (Herres & Gronholz, 1984)). The classic experimental reason is due to insufficient acquisition time combined with inadequate data processing, which results in the Fourier transformation of a truncated FID. In our analysis, ringing was the least commonly observed artifact and was only observed in IC-MS spectra. Under these conditions, there are limitations on FID acquisition time and the separation of compounds by IC results in some scans observing only a small number of compounds with high relative concentrations. In discussions with the instrument vendor (ThermoFisher Scientific), ringing artifacts in our Orbitrap instruments are due to a rare failure in processing software to handle an edge case when the minimum acquisition frequency approaches the Nyquist frequency under a specific range of injection times. Thus, the edge case condition results in a software failure that mimics a classic Fourier transform truncation artifact.

Partial ringing is more difficult to explain; however, the observation of peaks that exhibit ringing in one scan and partial ringing in other scans indicate that the two phenomena are most likely related. Also, partial ringing itself rarely occurs next to an intense peak, but rather in relative vicinity of an intense peak. Both observations suggest that partial ringing may result from ringing suppression on the instrument. The exact mechanism by which ringing is suppressed remains unclear without access to the raw time domain data acquired by the instrument and more detail on the algorithms used to process and transform this data. From discussions with ThermoFisher Scientific, these artifacts may be experimentally due to amplitude modulation of the FID that could have an electronic or vibrational origin.

Scan-level inconsistency of fuzzy sites and the absence of chemical phenomena with explanatory m/z patterns strongly suggests an artifactual origin for fuzzy sites. We hypothesize that fuzzy sites are another form of Fourier transformation artifact; specifically, they are the mass spectrometry equivalent to Gibbs phenomena, where transformation of an FID containing a discontinuity results in the production of many oscillations in a small subdomain of the transformed space. These oscillations would result in many peaks in a small m/z window, which is consistent with the scan-level appearance of fuzzy sites. At first glance, this scan-level manifestation appears inconsistent with the pseudo-Gaussian appearance of fuzzy sites at the aggregate level. This discrepancy can be explained by the central limit theorem. As increasingly more scan-level fuzzy sites are aggregated together around an expected point or region, which is essentially equivalent to averaging multiple small distributions together, the aggregate will converge to a larger Gaussian-like distribution.

In discussions with ThermoFisher Scientific, background electronic signals from accidentally oscillating electronic components either inside the instrument or from outside sources could give rise to phenomena that we detect as fuzzy sites in our non-Lumos Fusion spectra. These signals would appear to “hop” around from scan to scan since they are fundamentally unstable with respect to frequency and amplitude. Furthermore, a range of experimental conditions could influence this hopping, including proximity, temperature and other atmospheric conditions, and sample ion complexity.

### Fuzzy sites reduce robustness of downstream data analyses

This hypothetical possibility of impacting data analysis is demonstrated in the following classification results (Table 1). In this example, spectra ranging from 150 to 1600 m/z were acquired from paired cancer and non-cancer lung tissue sample non-polar extracts in an approximately random order with respect to sample class (see Supplemental Figure 7). Random forest was then used to train a classifier that would classify the samples based on LipidSearch peak assignments to expected lipids. The resulting classifier effectively classifies the samples into the appropriate class; however, these classifiers make extensive use of peaks that are present within fuzzy sites. Although these classifications are accurate, the classification is heavily based on artifactual features without direct molecular interpretation. Also, these same issues would have significant impact on differential abundance analyses as well.

### Mitigating the effects of fuzzy sites on downstream data analyses

The fuzzy-site/sample-class confound can be mitigated by simply removing fuzzy site assignments from the assignment lists prior to training the classifiers. However, care must be taken when performing this operation as a sample-by-sample removal of these sample class specific features can encode artifactual class information into the peaklist. For example, if the cancer class of spectra has a fuzzy site from 500 to 502 m/z but not the non-cancer, removal of and therefore absence of peaks from 500 to 502 m/z is also cancer-specific and therefore artifactually usable by machine learning methods. One solution is to remove the union of the m/z regions of all fuzzy sites from all spectra in an experiment. This prevents the accidental absence encoding of class information into the data, but has the disadvantage of removing larger regions of spectrum as the number of samples and classes increases. Table 1 shows the effect of fuzzy site removal on feature importance in classification. Twenty random forests were trained on LipidSearch assignment features derived from Thermo Tribrid Fusion FT-MS spectra collected on cancer and non-cancer lung tissue samples (Sample D). The top features based on mean importance of each feature across the 20 random forests are reported. Features in red came from the HPD region of a fuzzy site in at least one sample (HPD features). Without HPD feature removal, on average 6 out of the top 30 features are HPD features. But as mentioned earlier, it is non-trivial to remove HPD features correctly. If a feature is within an HPD region in any spectrum, the proper action is to remove that feature from all spectra, else these features may still contribute to classification. In the latter case, the artificial absence of that feature in some spectra can falsely boost its importance in the spectra that still contain that feature ad indicated by the encoding removal results in Table 1B. But when HPD features are removed in a non-encoding manner, no HPD feature makes it into the top 30 mean importance list (Table 1C). In this example, perfect disambiguation of cancer and non-cancer was possible with and without proper HPD removal; however, proper HPD feature removal is crucial to ensuring that important features represent true biological variance between the sample classes. Additionally, this demonstrates that classification accuracy is not necessarily an indication of classifier quality or robustness, especially when artifacts are present.

## Conclusions

With our HPD detector and ringing detector tools, we have identified and characterized three distinct types of artifacts that produce large numbers of peaks: fuzzy sites, ringing and partial ringing. In some cases, these artifacts can account for roughly 1/3 of the peaks detected in a given spectrum but only a small portion of total m/z range. Ringing is a known Fourier transformation artifact and we hypothesize that partial ringing and fuzzy sites are Fourier transformation-based artifacts as well. Partial ringing appears similar to ringing but suppressed at the scan level, with the ringing pattern only visible with multiple scans aggregated. Fuzzy sites have a more distinct pseudo-Gaussian appearance and were only observed in spectra from non-Lumos Fusion Tribid instruments.

All three artifacts complicate assignment and confound experimental interpretation; however, our study focused primarily on the fuzzy sites as they differed significantly from known artifacts and were particularly problematic for our classification studies. The correlation between fuzzy site location and sample class increases the probability of class-specific misassignment, introducing sample-specific artifactual features. Ultimately the presence of these artifacts produces brittle classifiers and complicates the characterization of true biological variance between sample classes using mass spectrometry. The results presented in this study reflect both the severity of this artifact for untargeted experiments, but also the deficiency in assignment tools specifically designed for untargeted analyses. The methods and tools presented in this study detect and remove HPD-sites in a non-encoding manner (i.e. will not encode sample class into spectra in the condition that HPD location is class specific), while providing sufficient protection from fuzzy sites with existing assignment methods. Ultimately, better assignment pipelines will be necessary as experiments grow in scale. A spectral analysis approach that can leverage peak correspondence between scans to infer if a peak is artifactual or real would produce higher quality peaklists for downstream data analysis, assignment, and interpretation.

## Resource Sharing

Code and data used for this manuscript are available here: https://figshare.com/s/700ea5fde9c2229c1f9c.

## Acknowledgements

This work was supported in part by National Science Foundation grant NSF 1252893 (Hunter N.B. Moseley), National Institutes of Health grants NIH 1R03CA211835-01 (Chi Wang and Robert Flight), NIH 1R01ES022191-01 (Teresa W.-M. Fan, Richard M. Higashi, Hunter N.B. Moseley, and Michael Nantz) and NIH 1U24DK097215-01A1 (Richard M. Higashi, Teresa W.-M. Fan, Andrew N. Lane, and Hunter N.B. Moseley), and Grant-in-Aid award from the American Heart Association (AHA16GRNT31310020 to Q.J.W.). We would like to thank Wenzhu Zhang and Brian T. Chait in the National Resource for the Mass Spectrometric Analysis of Biological Macromolecule at the Rockefeller University, and Shruti Nayak, Avantika Dhabaria, and Beatrix Ueberheide at the Proteomics Resource Center at the New York University Langone Medical Center for providing us with spectra for our bioinformatics analyses. We would also like to thank Thomas Wilson and the High Resolution Metabolomics Laboratory (HRML) at the Institute of Biological, Environmental and Rural Sciences for Q-Exactive+ data. We would like to thank Timothy Fahrenholz for collection of direct infusion spectra on the Fusion. We would like to thank Qiushi Sun for collection of ICMS spectra on the Fusion. Finally, we would like to thank Mike Senko from ThermoFisher Scientific for the extremely helpful discussions on the known and possible origins of the artifacts we have observed in Orbitrap FT-MS instruments.

---

**Supplemental Figure 1:**
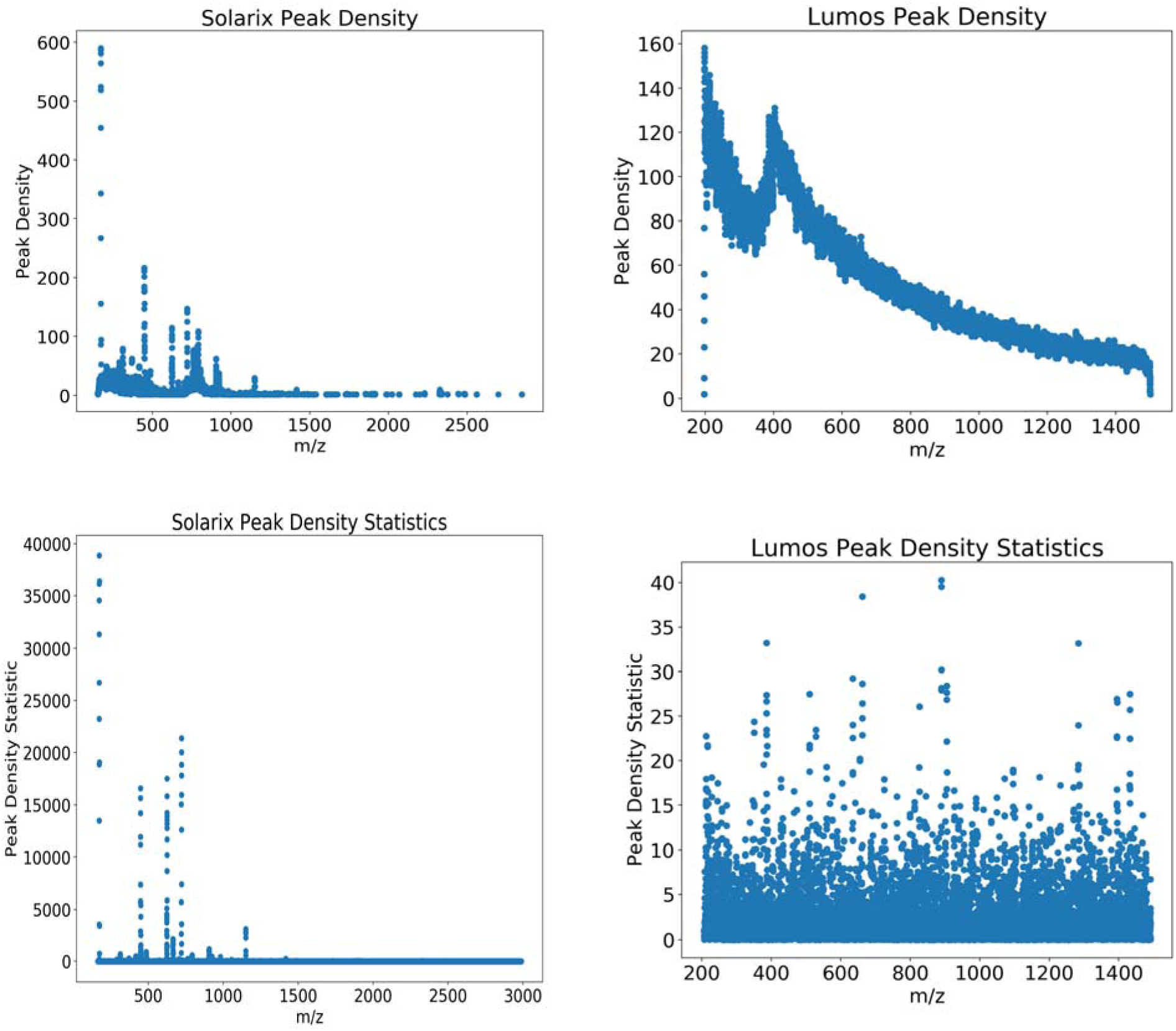
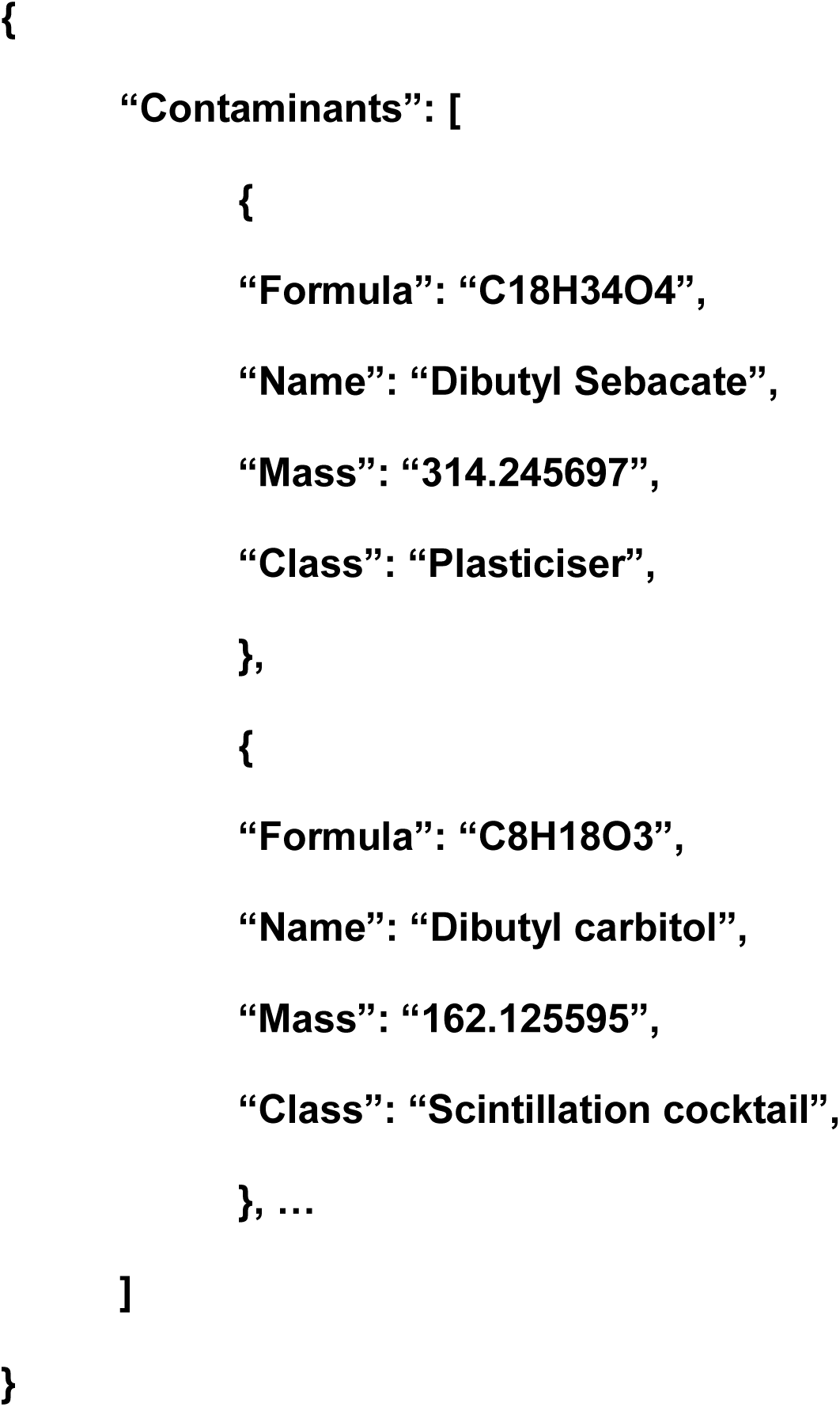
Additional Peak Density Examples. We have performed peak density analyses on both our Solarix ICR instrument and a Thermo Lumos Tribrid Fusion instrument, a more advanced version of the Tribrid Fusion. In the ICR, severe HPD phenomena are present due to ringing phenomena. It is not clear if the ringing artifacts from ICR are identical in origin to those in Orbitrap spectra. Our Lumos examples show no obvious HPD artifacts of any kind. In both the ICR and the Lumos, the peak density decreases with increasing peak density but not monotonically.

**Supplemental Figure 4:**
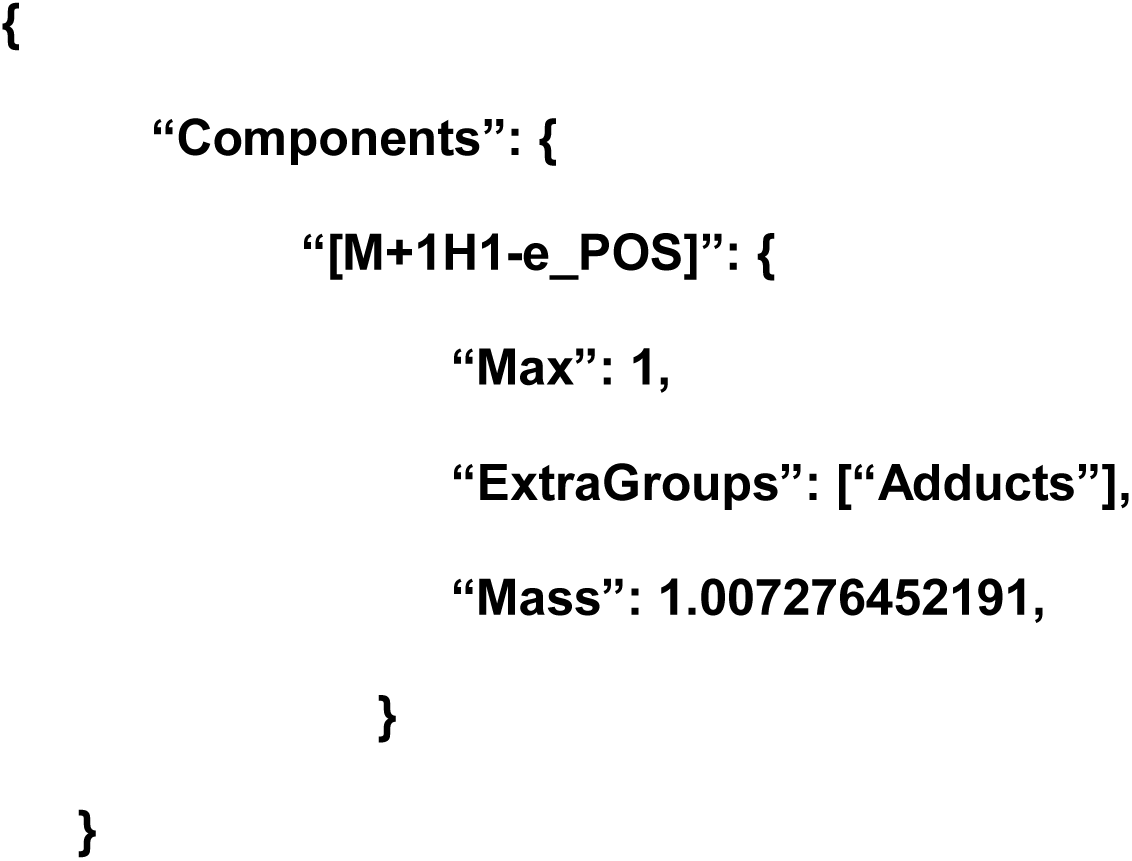
Adducts JSON.

**Supplemental Figure 5:**
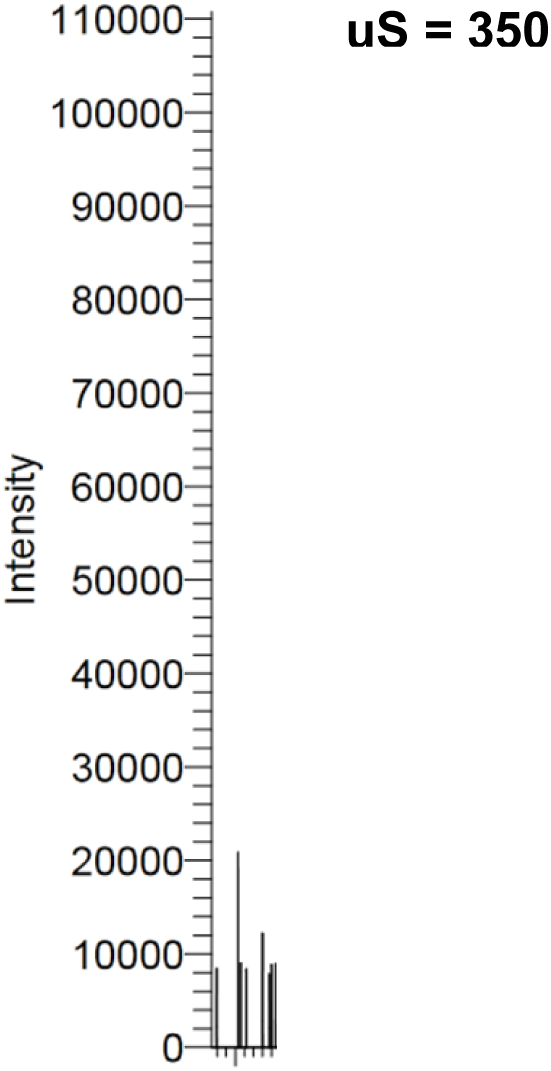
High uS Fuzzy Site. At very high microscan settings (Sample A), fuzzy sites become almost uniform and span a wide m/z window. Although still responsible for artifactual peaks, the significant intensity difference allows the distinguishing of true signal.

**Supplemental Figure 6:**
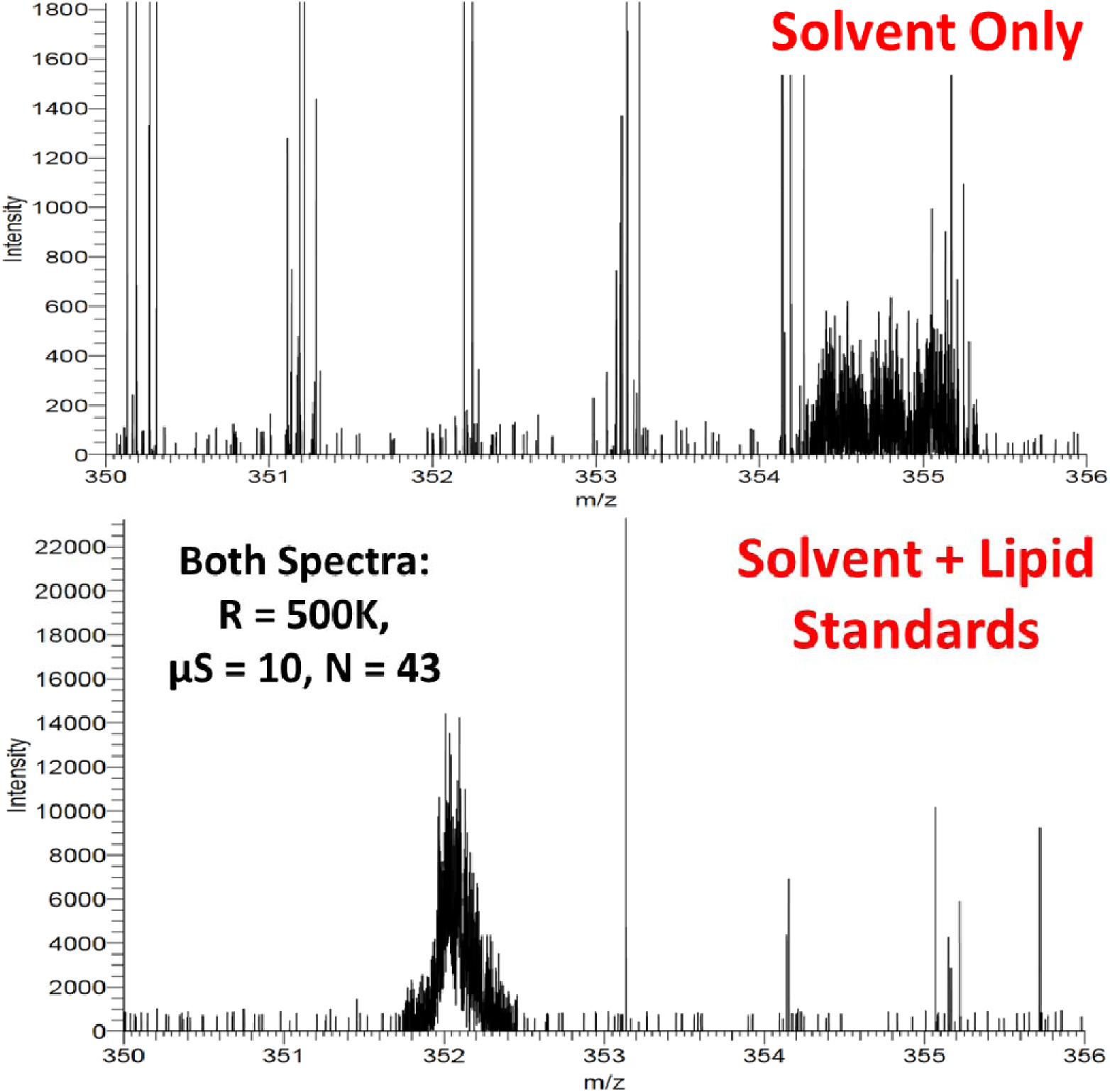
Fuzzy Sites vary with Sample Composition. A small change in chemical composition (Sample A with and without lipid standards) changes fuzzy site location. With only solvent, there is a fuzzy site at 354.8 m/z. With lipid standards, this fuzzy site shifts to 352.1 m/z. The number of fuzzy sites will remain constant, but will all be shifted by roughly the same m/z.

**Supplemental Figure 7:**
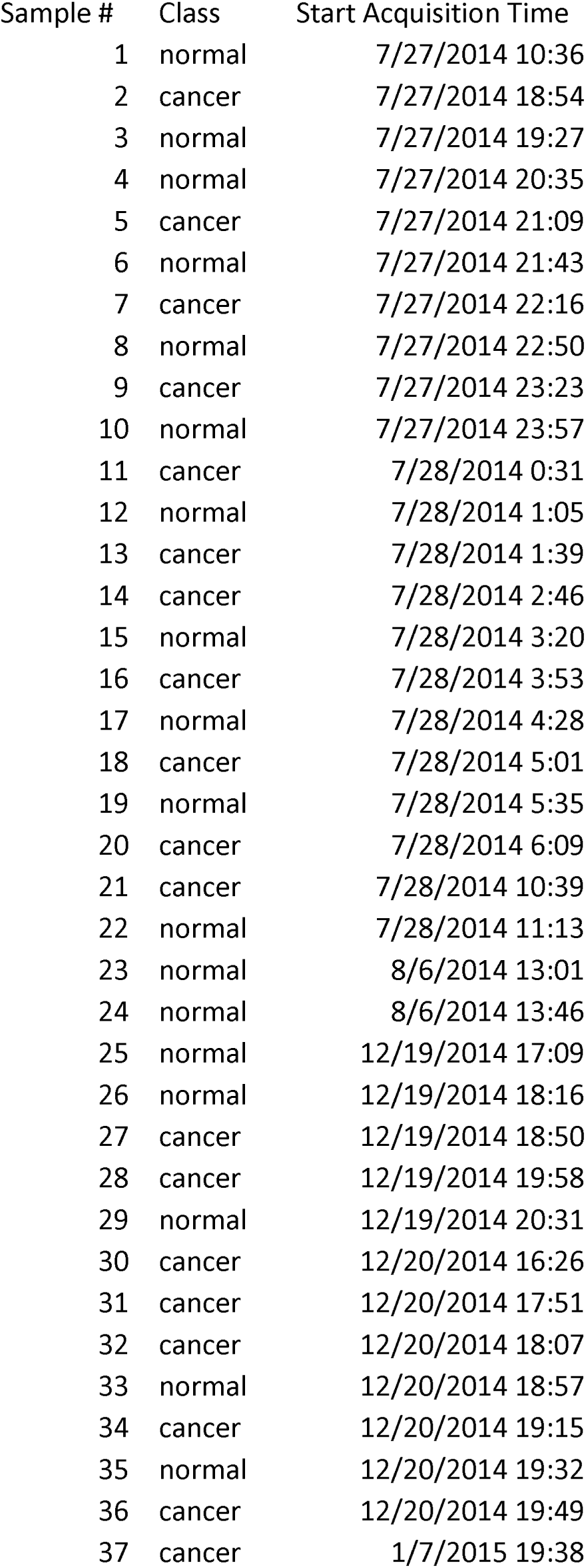
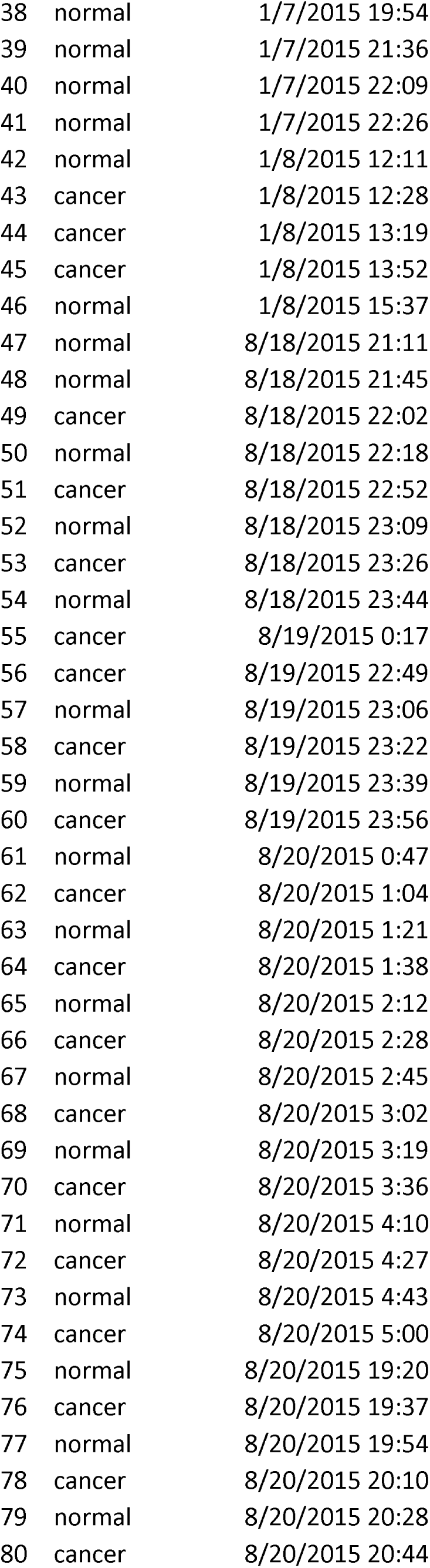
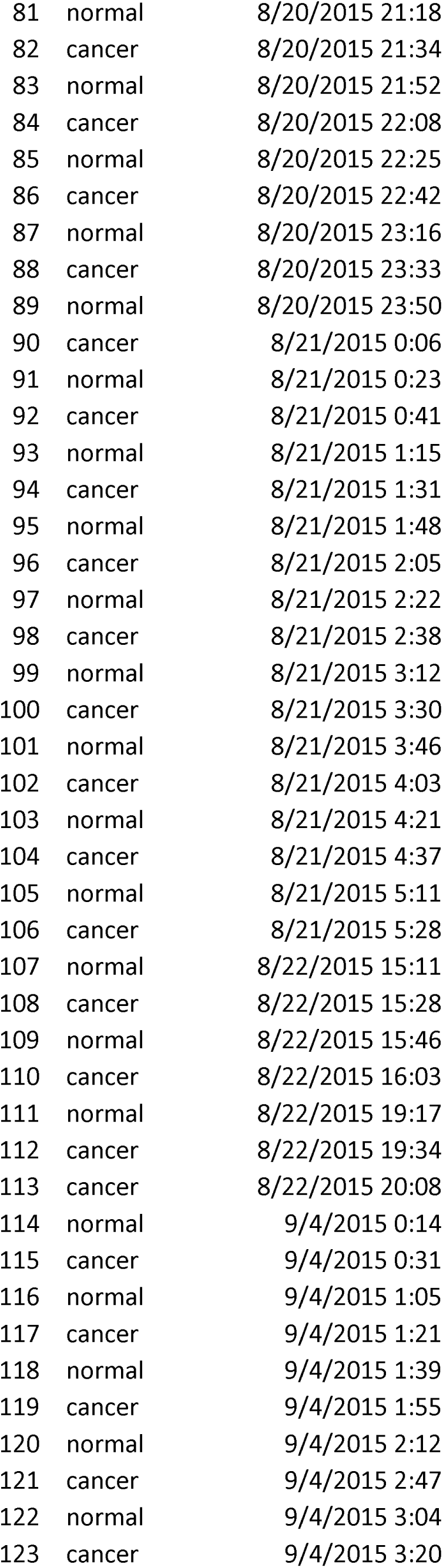
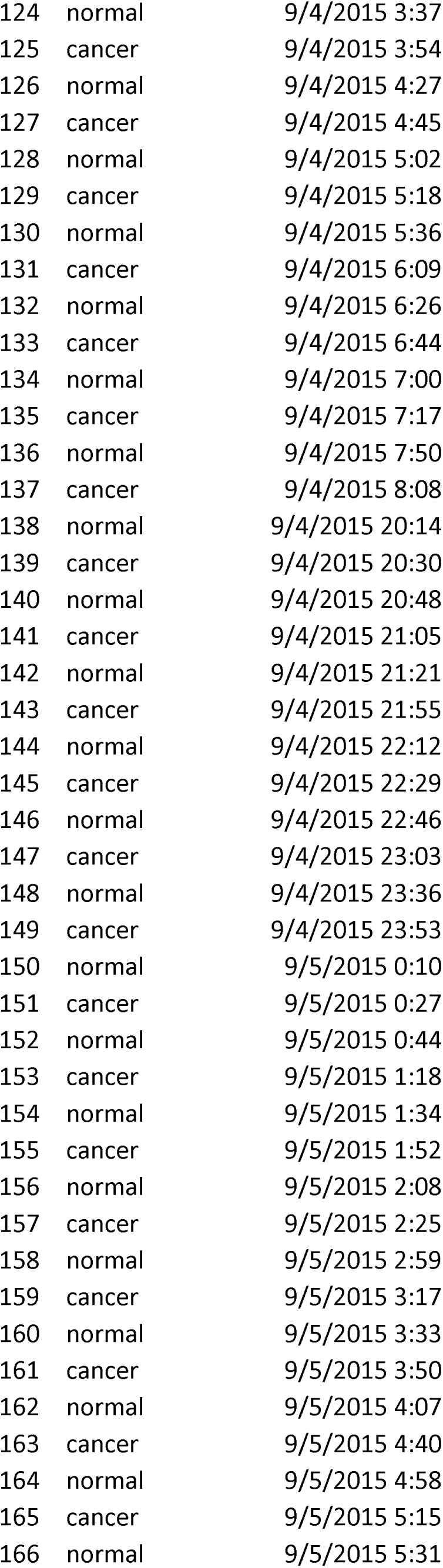
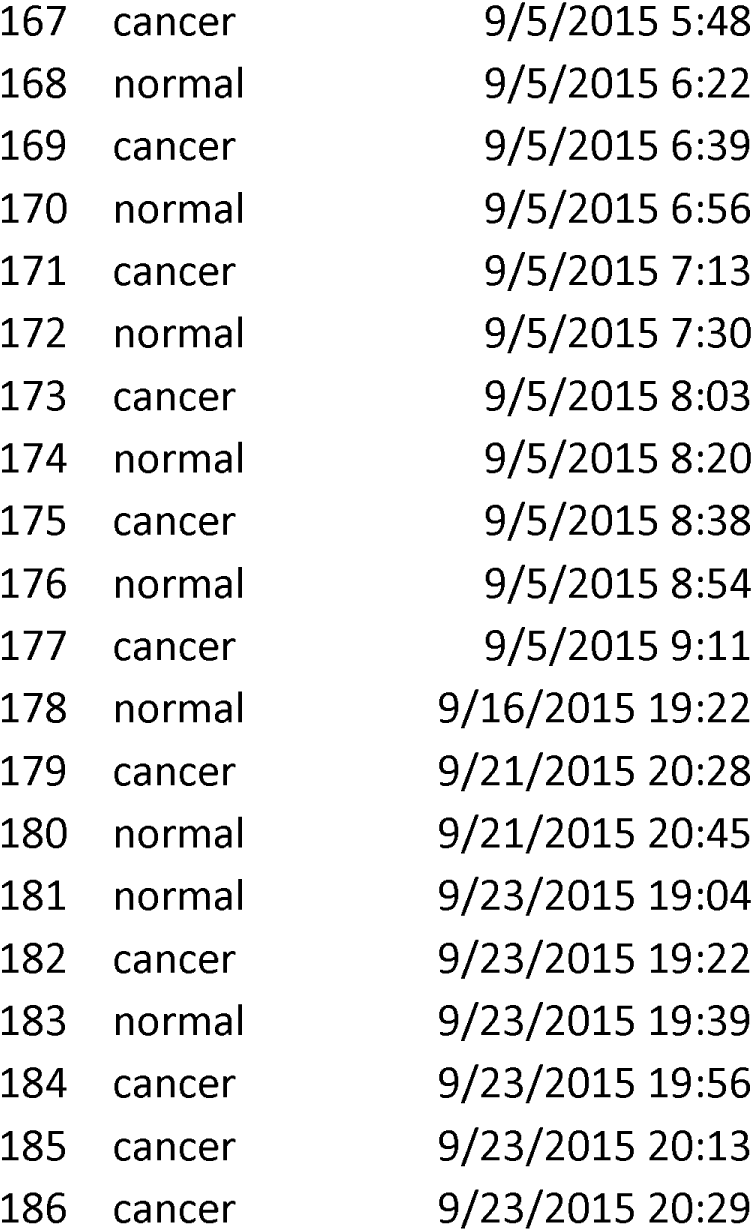
Spectral Acquisition Times for Sample D. Spectra for sample D show no significant “batches” with respect to time. Most samples were run sequentially over a time period from 7/2014 to 9/2015.

